# Creating wounds in cell monolayers using micro-jets

**DOI:** 10.1101/2021.01.07.425744

**Authors:** Cristian Soitu, Mirela Panea, Alfonso A. Castrejón-Pita, Peter R. Cook, Edmond J. Walsh

## Abstract

Many wound-healing assays are used in cell biology and biomedicine; they are often labor intensive and/or require specialized and costly equipment. We describe a contactless method to create wounds with any imaginable 2D pattern in cell monolayers using micro-jets of either media or an immiscible and biocompatible fluorocarbon (i.e., FC40). We also combine this with another method that allows automation and multiplexing using standard Petri dishes. A dish is filled with a thin film of media overlaid with FC40, and the two liquids reshaped into an array of microchambers in minutes. Each chamber in such a grid is isolated from others by fluid walls of FC40. Cells are now added, allowed to grow into a monolayer, and wounds created using the microjets; then, healing is monitored by microscopy. As arrays of chambers can be made using the media and Petri dishes familiar to biologists, and as dishes fit seamlessly into their incubators, microscopes, and workflows, we anticipate this assay will find wide application in biomedicine.

## Introduction

Wound-healing assays are widely used to investigate, for example, skin repair, angiogenesis, morphogenesis, and metastasis.^1-5^ Cell movements in and around wounds are complex, and governed by four major interlinked parameters: cell autonomous behaviours, cell-cell interactions, matrix compositions, and concentrations of critical solutes.^6^ These are often analysed *in vitro* by creating a wound – a cell-free area – in a confluent monolayer of cells (either by excluding cells from initially entering it, or removing cells from it), and then tracking cells as they repopulate the cleared area.^7^

A cell-free area can be created by surrounding it with some kind of removable barrier (e.g., in the form of a solid, liquid, or gel), but this can leave residues on the substrate that may alter cell behaviour during repopulation.^6^ Alternatively, the ‘scratch assay’ is commonly applied to clear cells from a selected part of a confluent monolayer, and its versatility, simplicity, and low cost make it popular.^8^ However, it is generally performed by hand – and so often lacks reproducibility (e.g., because application of too much force damages the substrate or underlying matrix), and – if automated – the associated machinery can be costly.

Wounds can also be created, for example, by electrical pulses,^9^ laser ablation,^10^ or laminar flows,^11^ but these approaches have not been widely adopted, possibly due to the requirement for specialised equipment.

Here, we introduce a contactless method to create wounds in 2D monolayers using micro-jets. Stationary jets of media have been used previously to detach cells, and so quantify the strength of cell adhesion.^12-13^ We attach an aqueous or fluorocarbon micro-jet to a 3-axis traverse, and so create wounds with almost any 2D geometry in seconds by jetting cell-media or an immiscible and biocompatible fluorocarbon (FC40) onto a monolayer of cells. We then combine this with “freestyle fluidics”^14-17^ to multiplex wound production. Arrays of hundreds of chambers–each containing a wound in a monolayer of cells–can be constructed in minutes on Petri dishes, which gives potential to screen drugs that might affect healing. As wounds with virtually any shape, size or scale can be made using media and dishes familiar to biologists, we anticipate this assay will find wide application in biomedicine.

## Results

### Principle of wounding

**Figure 1** illustrates two ways of creating wounds with micro-jets in confluent monolayers of mammalian cells. In the first (**Fig 1Ai**), the nozzle of a stainless-steel needle (held by a 3-way traverse, connected to a syringe pump, and filled with media) is lowered until 0.4 mm above the cells. When it starts jetting media, the resulting shear stress dislodges attached cells (**Fig S1A**).^13^ Moving the nozzle along a linear path then creates a linear wound filled with media. In practice, wounds are generated easily and quickly in monolayers of mouse C2C12 cells, with wound width being varied by overlapping jetting lines (**Fig. 1A**ii).

**Figure 1.**
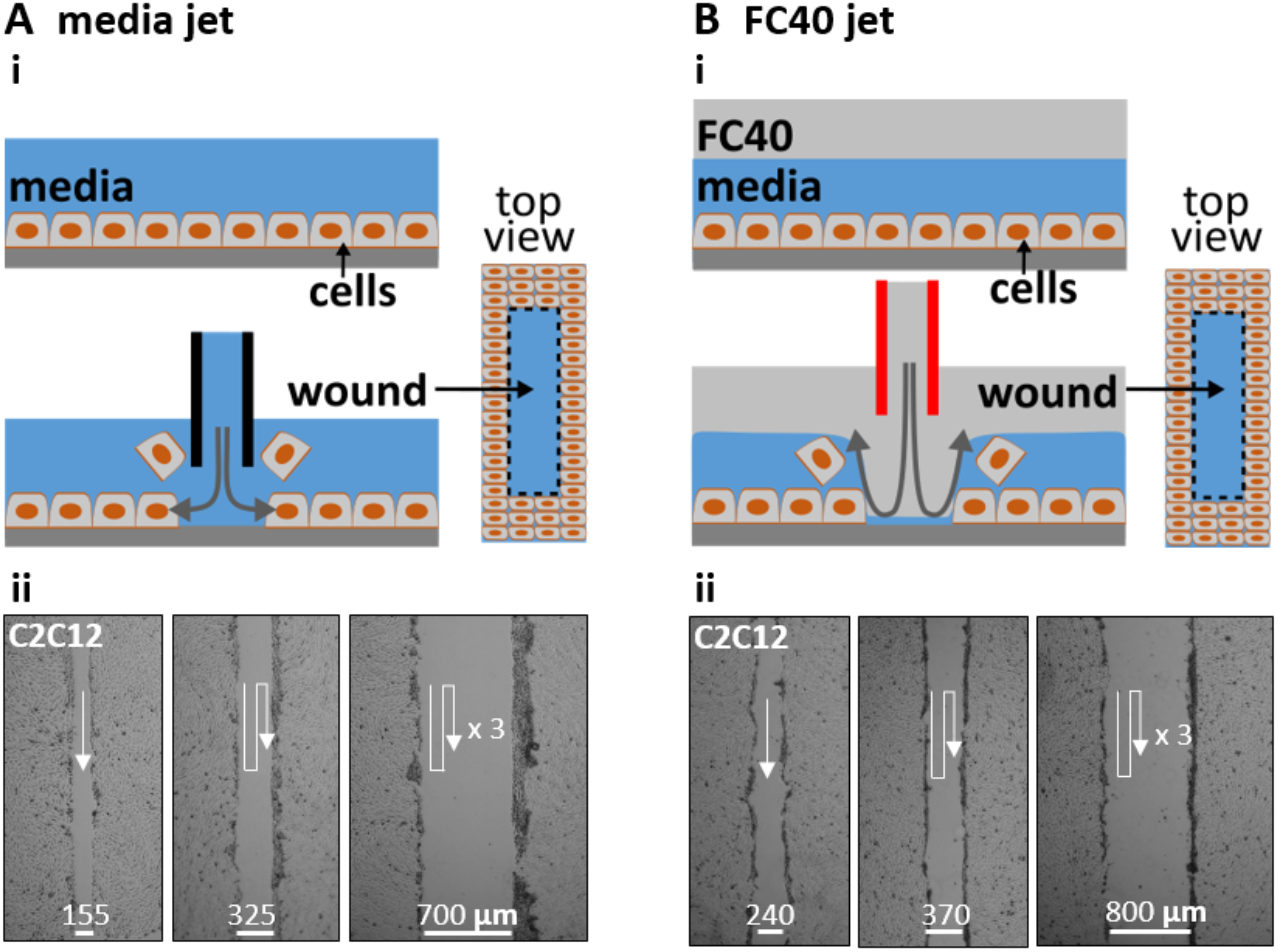
Creating wounds in a monolayer of mouse C2C12 myoblasts. **A**. With media jet. (**i**) Cartoon showing a monolayer before and after the nozzle of a steel needle filled with media is lowered until just above the cells; on jetting media, jet momentum detaches some cells from the surface, and moving the nozzle along a predefined path (here into the page) creates a wound (as shown in the top view). (**ii**) Images of wounds of different widths produced by jetting overlapping lines (white arrows: jetting paths). **B**. With FC40 jet. (**i**) Cartoon showing a monolayer overlaid with FC40 before and after the nozzle of a steel needle filled with FC40 is lowered into the FC40 overlay until just above the media; on jetting FC40, jet momentum forces the FC40:media interface down so it plays on the monolayer to dislodge cells, and moving the nozzle again creates a wound. (**ii**) Images of wounds with different widths created by jetting overlapping lines (white arrows: jetting paths).

In the second method (**Fig. 1B**), the monolayer and media are first covered by a layer of immiscible FC40. Although media is less dense than this overlay and one might expect buoyancy to drive it above the FC40, media remains firmly attached to the polystyrene dish held by stronger interfacial forces. This fluorocarbon is optically transparent, and freely permeable to O_2_ and CO_2_ so cells can be grown under it as usual in a CO_2_ incubator. The nozzle of a steel needle filled with FC40 is lowered through the fluorocarbon until just above the surface of the media, and the nozzle now jets FC40 instead of media (**Fig 1Bi**). Then, the momentum of the immiscible FC40 push media aside to detach cells, and – once the nozzle has passed – media refills the wound (**Fig S1B**). As before, wound width can be varied by overlapping jetting lines (**Fig. 1Bii)**. In this case, the bottom of the dish is pre-wetted with media (and initially covered with cells and/or materials derived from cells and serum), and jet momentum is insufficient to sweep all the aqueous phase (and cell material) from the substrate. Consequently, jetted FC40 does not adhere to the bottom; instead, it remains in the fluorocarbon overlay.

### Parameters affecting wound dimensions

Experimentalists often study a number of linear wounds of the same length and width. As wound length is easily controlled by starting and stopping the pump at appropriate times or places, we analyzed other parameters affecting wound width in detail, including nozzle diameter (*D*_*nozzle*_), height of the nozzle above the substrate (*H*), volumetric flowrate 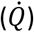,and speed of the traverse (*V*_*traverse*_) when making wounds (length 2.5 mm) in C2C12 myoblasts.

At a low media flowrate of 7.5 µl/s, jet momentum is insufficient to clear all cells off the substrate in the jetting line; we call this ‘failure’ (**Fig. 2Ai**, left). Increasing flow then produces progressively wider wounds (**Fig. 2Ai**, right). With an FC40 jet, failure occurs at a lower flowrate of ∼5.4 µl/s, and increasing flows again produce wider wounds (**Fig 2Aii**). Quantitative analysis confirms these general trends: failure occurs at low flows, increasing flows give wider wounds, and FC40 has a lower failure point and gives narrower wounds (**Fig. 2Aiii**). Increasing the traverse rate also leads to ‘failure’ and reduction in wound width with both media and FC40 jets (**Fig. 2B**).

**Figure 2.**
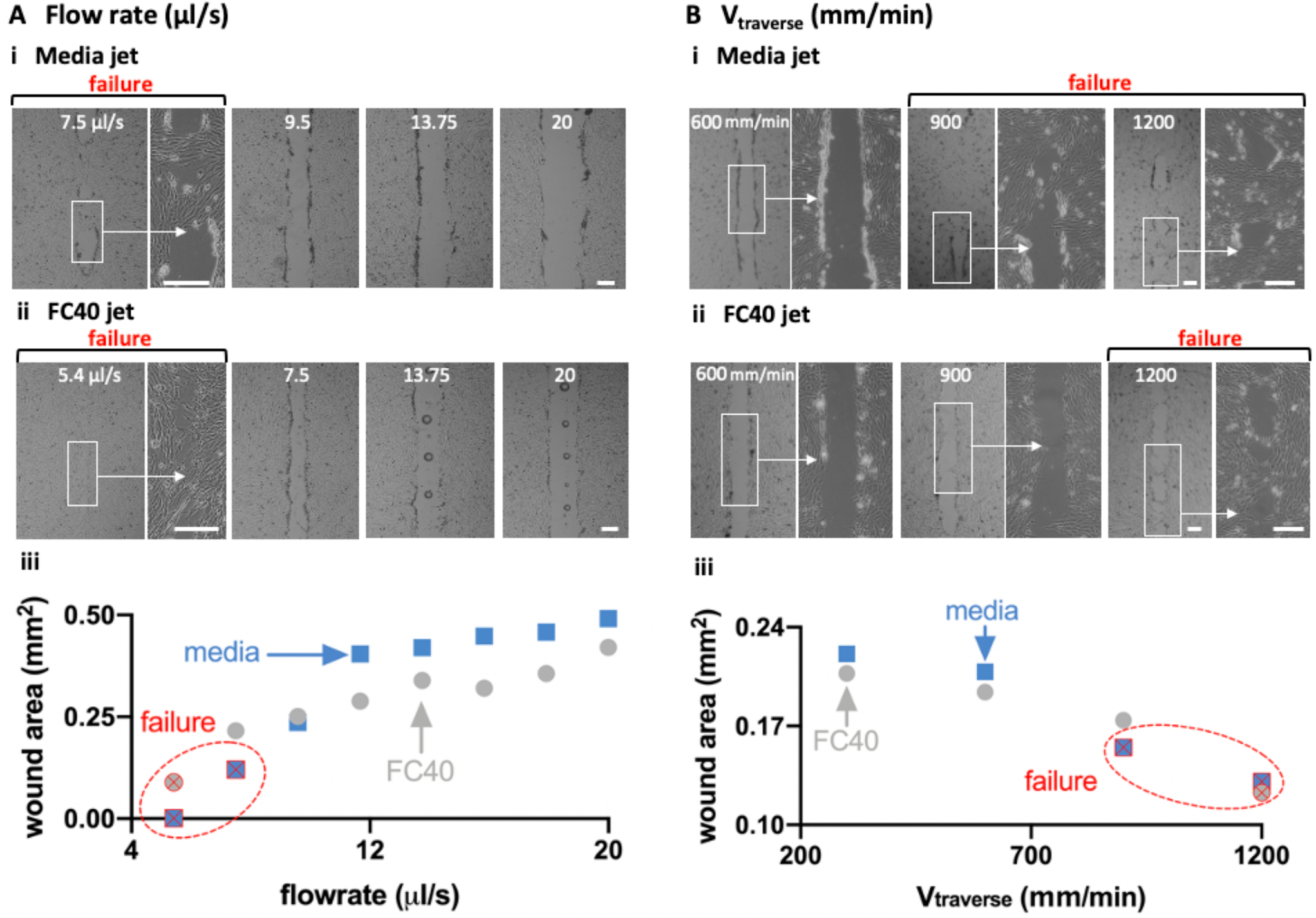
Effects of varying flowrate 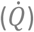 and traverse speed (*V*_*traverse*_) on wound width and area (*D*_*nozzle*_ = 60 μm, *H* = 0.4 μm, wound length = 2.5 mm). Scale bars: 200 µm. **A**. Varying 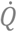 as *V*_*traverse*_ is held constant at 300 mm/min. (**i**) For media jets, wound failure occurs at 7.5 µl/s, and increasing flow produces wider wounds. (**ii**) For FC40 jets, wound failure occurs at 5.4 µl/s, and increasing flows again yield wider wounds (often with a central column of FC40 drops that adhere to the dish). (**iii**) Relationship between (cell-free) wound area and flowrate. **B**. Varying *V*_*traverse*_ as 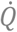 is held constant at 8 µl/s. (**i**) For a media jet, increasing the traverse rate leads to wound failure as the dwell time over a given area falls. (**ii**) For an FC40 jet, failure occurs at a higher traverse rate. (**iii**) Relationship between the cell-free wound area and traverse rate.

Flow profiles resulting as a jet emerges from a submerged and stationary nozzle to impinge on a flat substrate are complex.^18^ In our case, the substrate is initially covered with cells that are removed by jet momentum and shearing forces by the passing jet. The trends described above (**Fig. 2A**) are consistent with jet momentum playing a critical role. For example, the longer a jet spends above a specific point, the more persistent are the shearing forces so that more cells are dislodged (resulting in wider wounds). Moreover, when jet momentum falls below a critical value, wound production fails. In addition, FC40 is ∼1.8 times denser than media so it gives narrower wounds, with failure occurring at a lower flowrate and higher traverse rate.

Note that passage of an FC40 jet not only clears cells from the target area, but also leaves a column of FC40 droplets down the centerline of the wound (**Fig. 2Aii**; Soitu *et al*., 2020,^17^ describe a related phenomenon). This presumably arises as follows. At low flowrates, the FC40 jet cannot completely displace all media from the substrate, and FC40 never contacts polystyrene; then – as we have seen – media refills the wounded area, and jetted FC40 rejoins the overlay as it minimizes its interfacial area with the aqueous phase. However, as flowrate increases, momentum becomes sufficient to sweep media from some local areas of the substrate. Now, FC40 can contact local areas of the dish to remain stuck there as it preferentially wets polystyrene compared to media (and so gives a column of drops). These results show that linear wounds with chosen lengths and widths can easily be made by jetting and even shapes.

Assay geometry is another important parameter affecting migration into a wound.^19^ Wounds with almost any imaginable 2D pattern are easily generated by altering the path followed by the micro-jet – for example, from one straight or circular line (**Fig. 3A,B**), through many straight lines in a grid (**Fig. 3C**), to two curved lines plus some straight ones (which in the upper panel of **Fig. 3D** represent the two backbone strands and base-pairs in a double helix of DNA).

**Figure 3.**
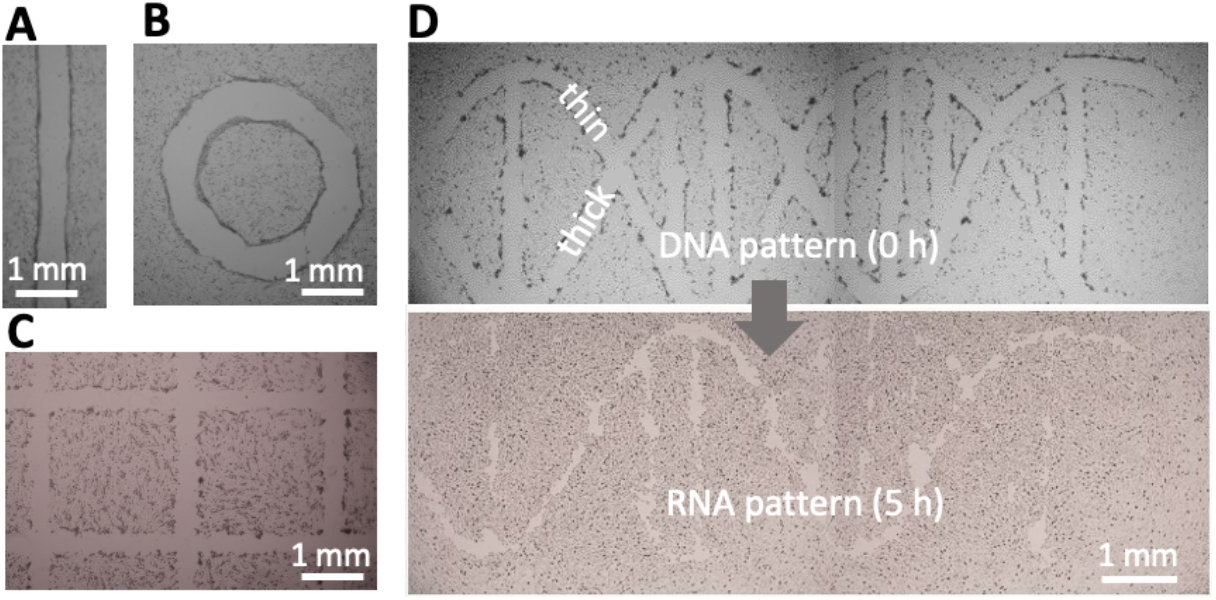
Wounding patterns. All shapes were created in 6 cm Petri dishes using media jets. **A**. Line. **B**. Circle.**C**. A complex wound (part of 16×16 grid). **D**. A wound shaped like a double helix of DNA (with one thick backbone strand + attached base, and one thin strand + base). After incubation (5 h), the thin strand has almost completely healed, so the structure looks like a single strand of RNA.

### Wound healing

We next illustrate healing of the wound shaped like a double helix where one helical ‘strand’ is initially thicker than the other (**Fig. 3D**, top). On incubation at 37^°^C, cells repopulate cleared areas to ‘heal’ the wound. As the thin strand heals quicker than the thick one, this leaves a ‘single RNA strand’ (**Fig. 3D**, bottom). The simpler wounds illustrated in **Figure 3A-C** also heal in <24 h (**Fig. S2**). These results show that wounds produced by jetting heal as expected.

We illustrate multiplexed wound healing using a ‘grid’ – an array of 256 microfluidic chambers in a 6 cm dish (**Fig. 4A**).^15^ Each chamber has a square footprint of 1.9 x 1.9 mm, holds <1 μl, and is isolated from others by fluid walls and ceilings of FC40. Such chambers are used much like wells in microtiter plates, albeit with one-hundredth the volume; liquids are added or removed simply by pipetting through FC40 instead of air. Then, fluid walls morph above an unchanging footprint within limits imposed by the advancing and receding contact angles around the footprint. When too much volume is added, the advancing contact angle is exceeded, the footprint expands, and neighboring chambers merge. Even so, the range of volumes that can be held before footprint expansion is greater than that held by a well in a 96-well plate. Unfortunately, the maximum volume that can be accommodated is less than the 2-6 μl needed to form a long linear wound by jetting media; therefore, an FC40 jet is used.

**Figure 4.**
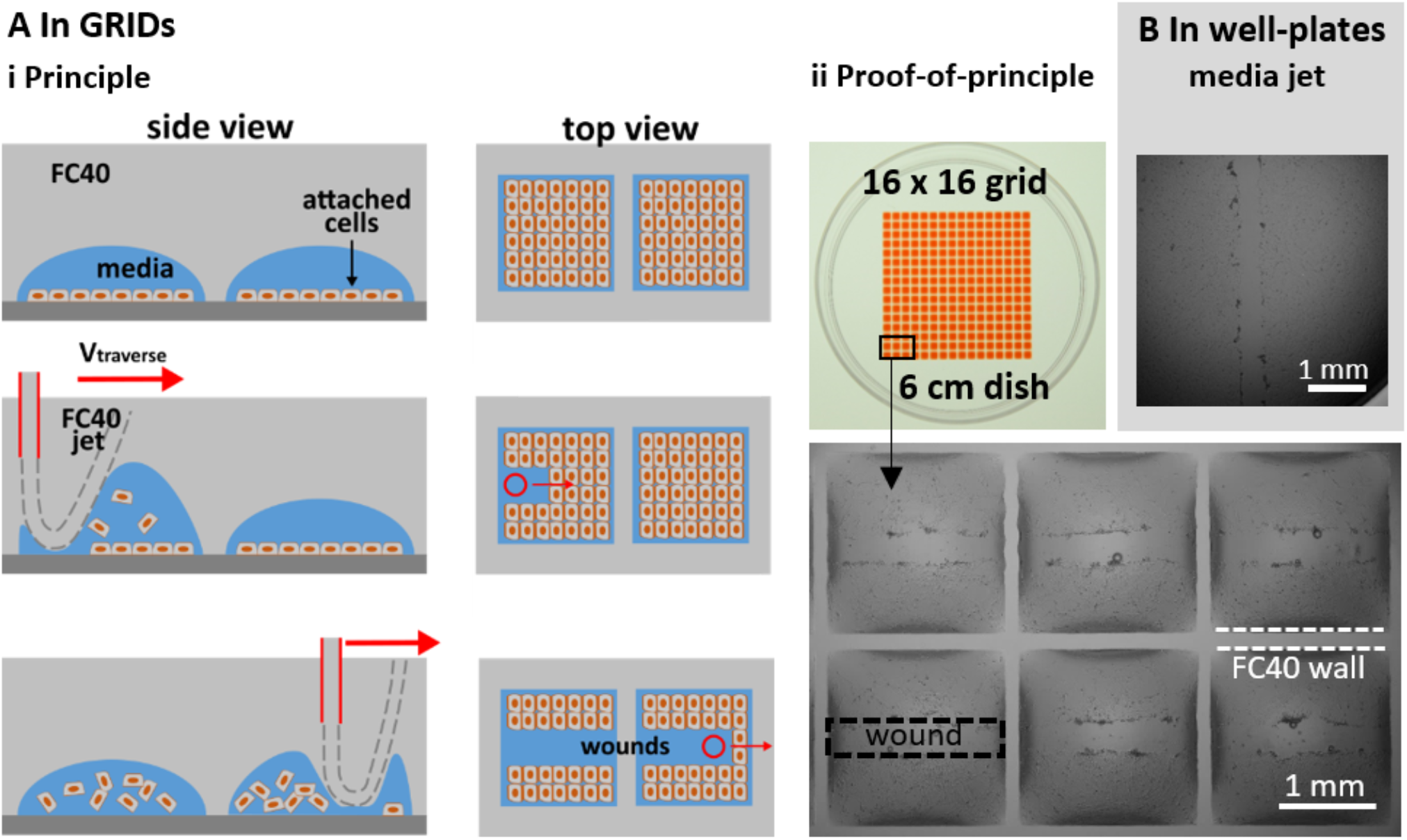
Multiplexed wounding. **A**. In grids, with FC40 jet. (**i**) Principle. Cells are seeded in chambers in a grid, grown to confluency, and the jet traverses above the centerline of each row; as it passes, the jet forces FC40;media interfaces down so they play on monolayers to detach cells. (**ii**) Image of grid. Chambers initially held 100 nl media, and 400 nl red dye is added to aid visualization. Zoom: images of wounds in 6 chambers. **B**. In well of a conventional 96-well plate, with media jet.

**Figure 4Ai** illustrates the principle. C2C12 myoblasts are seeded in each chamber, grown until confluent, and then the FC40 jet traverses across the middle of a chamber; the jet transiently forces the FC40:media interface down so it plays on the monolayer to dislodge cells, before the added FC40 returns to join the bulk FC40 in the overlay. Imaging then reveals the resulting wounds in the middle of each chamber (**Fig. 4Aii**).

Wounding can also be multiplexed using conventional 96-well plates and a media jet – as these can hold larger volumes (**Fig 4B**). Note the large black ring close to the well wall (**Fig. 4B**). This ‘edge effect’ prevents a clear view of much of the bottom of the well, and is due to the solid surrounding wall obscuring the view. In contrast, edge effects in chambers with fluid walls are much less (**Fig 4Aii**), and can be essentially eliminated using smaller volumes.^20^ These results show that our wound-healing assays can easily be multiplexed and used to screen drugs to see how they affect repopulation.

## Discussion

We describe a general contactless method to create wounds in a cell monolayer; micro-jets of media or an immiscible liquid (FC40) are projected on to the monolayer to detach cells (**Fig. 1, Fig. S1**). We establish working conditions and failure parameters for one cell type (mouse C2C12 myoblasts) and describe ways to alter wound dimensions (**Fig 2**). Wounds can have almost any imaginable 2D pattern (**Fig 3**), they heal as expected (**Fig. 3D, Fig S2**), and the approach can be multiplexed (e.g., using either an FC40 jet and a 6 cm dish with 256 chambers, or a media jet and a 96-well plate; **Fig. 4**).

Some limitations of the technology include the following. (i) A 3-axis traverse and syringe pump are essential (to produce and control the required jet). (ii) Whilst the method is a general one, conditions have to be tuned to produce wounds with specified widths in monolayers of different cell density and type, as both degree of confluency and cell type affect wound production. (iii) When using an FC40 jet, occasional FC40 drops are left behind in the wound, and these can potentially act as a physical barrier that prevents migration of cells into all areas of the wound (**Fig. 2A**).

In summary, we have developed a contactless method to produce wounds *in vitro*. This enables experimentalists to vary the size and shape of wounds with ease. The versatility of the methods also allow it to be tuned to accommodate samples with different cell-matrix characteristics, enhancing consistency and reproducibility over multiple experiments.

## Materials and methods

### General reagents

All reagents and materials were purchased from Sigma Aldrich (St. Louis, Missouri), unless otherwise stated. FC40^STAR^ (iotaSciences Ltd, Oxfordshire, UK) is FC40 treated using a proprietary method that improves fluid wall formation by jetting, and throughout we use the term ‘FC40’ to refer to it. In **Figure 4A**, allura Red – a water-soluble dye – was added to each chamber to aid visualisation.

### Cells

C2C12, an immortalized mouse myoblast cell line,^21^ was cultured in DMEM + 15% FBS. Prior to wounding, cells were plated in 60 mm tissue-culture-treated dishes (Corning, Merck product 430166) at a density of 10^6^ cells/dish, and grown for ∼48 h. For **Figure 4B**, cells were plated on 96-well plates (Corning, product 3596) at a density of 15,000 cells/well, and used for wounding ∼48 h later.

### Wounding and printing of grids

All wounds were made using custom-written software and modified ‘isoCell’ (used with 6 cm dishes) or ‘Pro’ printers (used with 96-well plates; **Fig. 4B**) provided by iotaSciences Ltd. Each printer consists of a 3-axis traverse, which moves two stainless-steel dispensing needles (Adhesive Dispensing Ltd, Milton Keynes, UK), one with a nozzle used for jetting media or FC40 (60 µm ID, 0.5 mm OD), and the other to add/remove media to/from chambers (250 µm ID, 0.5 mm OD). The two needles are connected to syringe pumps which control the jetting/dispensing of media/FC40.

Grids (**Fig. 4A**) were made with DMEM plus 10% FBS. To make a grid, 1 ml media is added to a 6 cm dish to cover the entire surface, and ∼0.9 ml removed to leave a thin film. This is now covered with 2 ml FC40, and the grid created as described by Soitu *et al*. (2018).^15^ Unless otherwise stated, wounds were created using *D*_*nozzle*_ = 60 µm, *H* = 0.4 mm,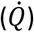, and *V*_*traverse*_ = 300 mm/min.

The DNA-shaped wound in **Figure 3D** was constructed after drawing the pattern in Inkscape (inkscape.org) followed by conversion to G-code, the programming language used by the printer.

### Imaging

All images of wounds were collected using a digital SLR camera (Nikon D7100 DSLR) connected to an epi-fluorescence microscope (Olympus IX53; 1.25X, 4X, 10X, 25X objectives) with translation stage and overhead illuminator (Olympus IX3 with filters). Brightness and contrast was enhanced by 20% for these images, for better visualisation of wounds. The image of the grid in **Figure 4B** was taken using a digital SLR camera (Nikon D610). Images of wounds in **Figure 2** were analysed using ImageJ (Rasband, 1997-2018) and areas of wounds extracted.

### Statistical analyses

Statistical analyses were performed using GraphPad Prism (San Diego, CA).

## Acknowledgements

We thank David Beeson’s group for providing the C2C12 cells. This work was supported by iotaSciences Ltd (who provided scholarships for C.S.), and a Royal Society University Research Fellowship (A.A.C.-P.), the Impact Acceleration Account of the Biotechnology and Biological Sciences Research Council (P.R.C. and E.J.W.), and awards from the Medical Research Council under the Confidence in Concept scheme (MC_PC_15029 to P.R.C. and E.J.W).

## Author contributions

C.S., P.R.C., and E.J.W. designed research, C.S., M.P., and E.J.W. performed experiments, and all authors wrote the paper.

## Data availability statement

The data that support the findings of this study are available from the corresponding author upon reasonable request.

---

**Figure S1.**
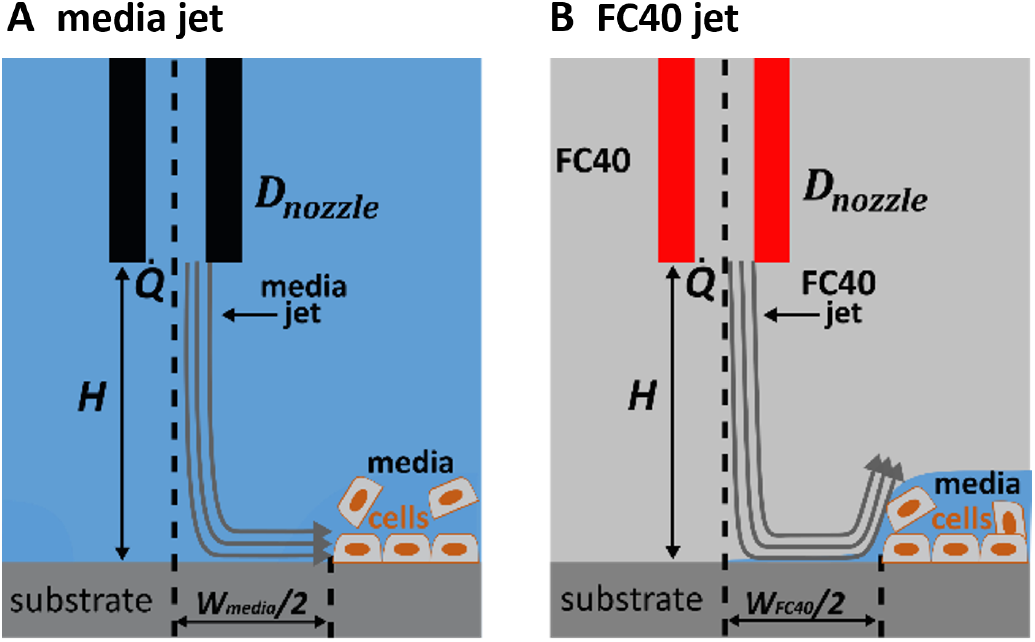
Parameters affecting wound width. Cells are detached by shear produced by the media or FC jet; this creates a wound of width *W*_*media*_. Unless otherwise stated, all experiments were performed using *D*_*nozzle*_ (nozzle diameter) = 60 μm, 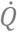 (volumetric flowrate) = 8 μl/s, *H* (height of nozzle above substrate) = 0.4 μm, *V*_*traverse*_ (traverse speed) = 300 mm/min. Streamlines of jets (grey) are for illustrative purposes only. **A**. Nozzle in media. **B**. Nozzle in FC40. Here, the FC40 jet displaces cells from the substrate but does not remove all the aqueous phase from the substrate under the nozzle, and – as the jet moves on – media flows back to refill the ‘wound’ area.

**Figure S2.**
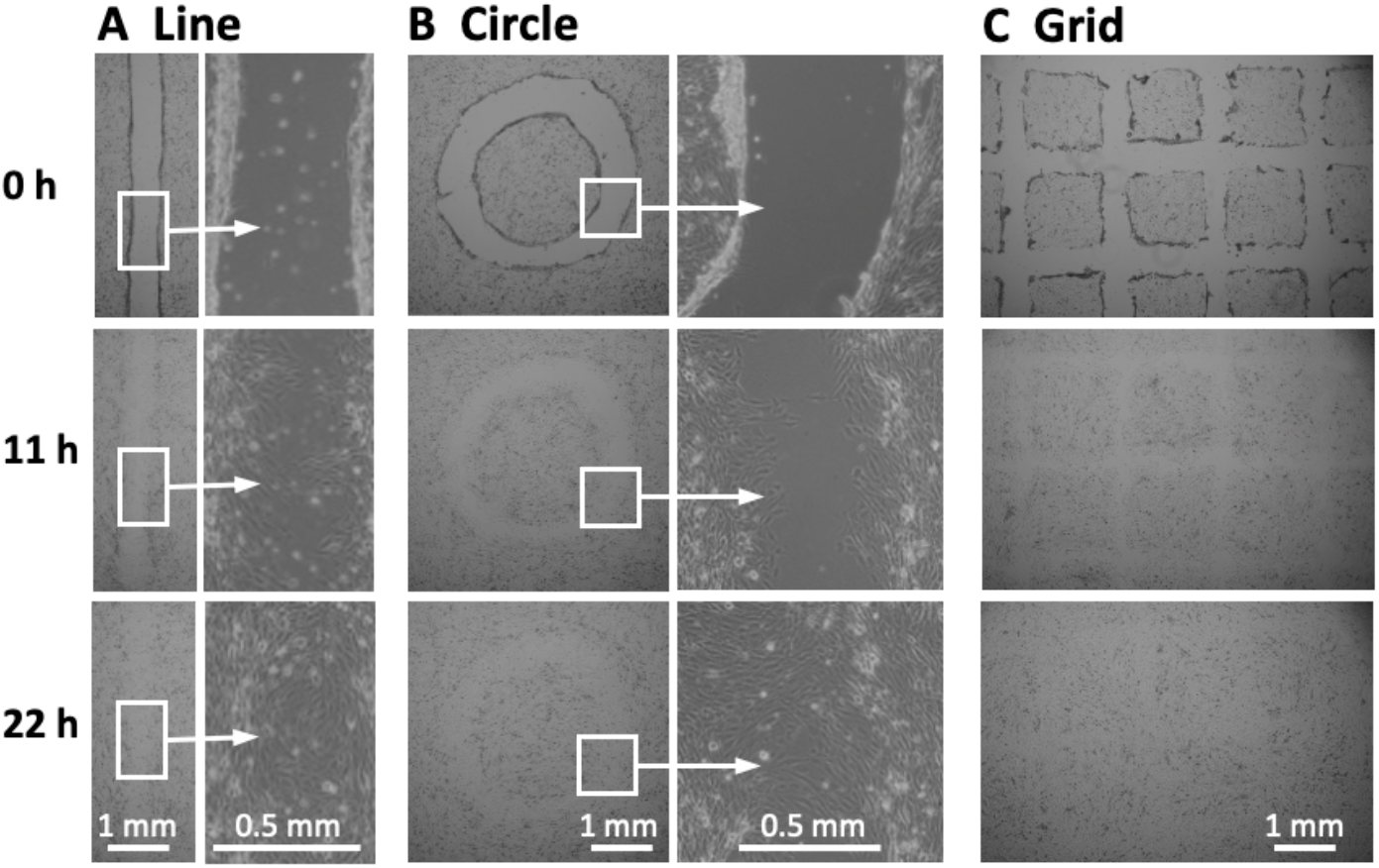
Healing of wounds similar to those shown in **Figure 3**. Images were collected at the times indicated. In all cases wounded areas are repopulated with cells after 22 h, and the different wounds yield subtle differences in the final structure of the cell sheet. **A**. Line. **B**. Circle. **C**. Grid.

## References

1. P. Friedl, D. Gilmour, Collective cell migration in morphogenesis, regeneration and cancer. Nat Rev Mol Cell Biol 10, 445–457 (2009).

2. Li L, He Y, Zhao M, Jiang J. Collective cell migration: Implications for wound healing and cancer invasion. Burn Trauma 1,21–6 (2013).

3. A. Grada, M. Otero-Vinas, F. Prieto-Castrillo, Z. Obagi, V. Falanga, Research techniques made simple: analysis of collective cell migration using the wound healing assay. J. Invest. Dermatology. 137, e11–e16 (2017).

4. M. Nakamura, A.N.M. Dominguez, J.R. Decker, A.J. Hull, J.M. Verboon, S.M. Parkhurst. Into the breach: how cells cope with wounds. Open Biol. 3,8 (2018);

5. D. Chouhan, N. Dey, N. Bhardwaj, B.B. Mandal, Emerging and innovative approaches for wound healing and skin regeneration: Current status and advances. Biomaterials 216:119267 (2019).

6. W.J. Ashby, A. Zijlstra, Established and novel methods of interrogating two-dimensional cell migration. Integrative Biology 4(11):1338–1350 (2012).

7. J.E. Jonkman, J.A. Cathcart, F. Xu, M.E. Bartolini, J.E. Amon, K.M. Stevens, P. Colarusso, An introduction to the wound healing assay using live-cell microscopy. Cell Adhesion & Migration 8(5):440–451 (2014).

8. C. Liang, A. Park, J. Guan, In vitro scratch assay: a convenient and inexpensive method for analysis of cell migration in vitro. Nature Protocols 2, 329–333 (2007).

9. C.R. Keese, J. Wegener, S.R. Walker, I. Giaever. Electrical wound-healing assay for cells in vitro. PNAS 101 (6) 1554–1559 (2004).

10. M.S. Hutson, Y. Tokutake, M.-S. Chang, J.W. Bloor, S. Venakides, D.P. Kiehart, G.S. Edwards, Science, 145– 149 (2003).

11. Y. Wei, F. Chen, T. Zhang, D. Chen, X. Jia, J. Wang, W. Guo, J. Chen, A Tubing-Free Microfluidic Wound Healing Assay Enabling the Quantification of Vascular Smooth Muscle Cell Migration. Scientific Reports 5, 14049 (2015).

12. A. Bouafsoun, A. Othmane, A. Kerkeni, N. Jaffrézic, L. Ponsonnet, Evaluation of endothelial cell adherence onto collagen and fibronectin: a comparison between jet impingement and flow chamber techniques. Materials Science and Engineering 26, 260–266 (2006).

13. W. Visser, M.V. Gielen, Z. Hao, S. Le Gac, D. Lohse, C. Sun, Quantifying cell adhesion through impingement of a controlled microjet. Biophysics Journal 108, 23–31 (2015).

14. E.J. Walsh, A. Feuerborn, J.H.R. Wheeler, A.N. Tan, W.M. Durham, K.R. Foster, P.R. Cook, Microfluidics with fluid walls. Nature Communications 8, 816 (2017).

15. C. Soitu, A. Feuerborn, A.N. Tann, H. Walker, P.A. Walsh, A.A. Castrejon-Pita, P.R. Cook, E.J. Walsh, Microfluidic chambers using fluid walls for cell biology. JPNAS 115, E5926–E5933 (2018).

16. C. Soitu, A. Feuerborn, C. Deroy A.A. Castrejon-Pita, P.R. Cook, E.J. Walsh, Raising Fluid Walls Around Living Cells. Science Advances. 2019, 5, eaav8002.

17. C. Soitu, N. Stovall-Kurtz, C. Deroy, A.A. Castrejon-Pita, P.R. Cook, E.J. Walsh. Jet-printing microfluidic devices on demand. Advanced Science 2001854 (2020).

18. M.D. Deshpande, R.N. Vaishnav, Submerged laminar jet impingement on a plane. Journal of Fluid Mechanics 114, 213–236 (1982).

19. K.K. Treloar, M.J. Simpson, D.L.S. McElwain, R.E. Baker, Are in vitro estimates of cell diffusivity and cell proliferation rate sensitive to assay geometry? Journal of Theoretical Biology 356, Pages 71–84 (2014).

20. C. Soitu, C. Deroy, A.A. Castrejón-Pita, P.R. Cook, E.J. Walsh, Using fluid walls for single-cell cloning provides assurance in monoclonality. SLAS Technology 25(3), 267–275 (2020).

21. J. Cossins, W.W. Liu, K. Belaya, S. Maxwell, M. Oldridge, T. Lester, S. Robb, D. Beeson, The spectrum of mutations that underlie the neuromuscular junction synaptopathy in DOK7 congenital myasthenic syndrome. Human Molecular Genetics 17, 3765–3775 (2012).

